# Real-time observation of flexible domain movements in Cas9

**DOI:** 10.1101/122069

**Authors:** Saki Osuka, Kazushi Isomura, Shohei Kajimoto, Tomotaka Komori, Hiroshi Nishimasu, Tomohiro Shima, Osamu Nureki, Sotaro Uemura

## Abstract

The CRISPR-associated protein Cas9 is a widely used genome editing tool that recognizes and cleaves target DNA through the assistance of a single-guide RNA (sgRNA). Structural studies have demonstrated the multi-domain architecture of Cas9 and sequential domain movements upon binding to the sgRNA and the target DNA. These studies also hinted at the flexibility between domains, but whether these flexible movements occur in solution is unclear. Here, we directly observed dynamic fluctuations of multiple Cas9 domains, using single-molecule FRET. The flexible domain movements allow Cas9 to adopt transient conformations beyond those captured in the crystal structures. Importantly, the HNH nuclease domain in Cas9 only accessed the DNA cleavage position during such flexible movements, suggesting the importance of this flexibility in the DNA cleavage process. Our FRET data also revealed the conformational flexibility of apo-Cas9, which may play a role in the assembly with the sgRNA. Collectively, our results highlight the potential role of domain fluctuations in driving Cas9-catalyzed DNA cleavage.

## INTRODUCTION

CRISPR (clustered regularly interspaced short palindromic repeats)-Cas (CRISPR-associated) systems were originally found as adaptive immunity systems against viruses and plasmids in bacteria and archaea (Jansen *et al*, 2002; Soria *et al*, 2005; Bolotin *et al*, 2005; Pourcel *et al*, 2005; Barrangou *et al*, 2007). Unlike other CRISPR-Cas systems that employ ensembles of Cas proteins to recognize and cleave nucleic acids, the type II CRISPR-Cas system utilizes the single RNA-guided endonuclease Cas9 protein for the destruction of foreign nucleic acids (Shmakov *et al*, 2017). *Streptococcus pyogenes* Cas9 (henceforth, Cas9) has been widely used as a powerful genome editing tool (Jinek *et al*, 2012; Mali *et al*, 2013; Cong *et al*, 2013), especially since Cas9 can be programmed by a synthetic single-guide RNA (sgRNA) to cleave any specific DNA sequence followed by a protospacer-adjacent motif (PAM) (Jinek *et al*, 2012). In addition, Cas9 has been applied to visualize, modify and express endogenous target genes (Hsu *et al*, 2014; Terns & Terns, 2014; Konermann *et al*, 2014; Sternberg & Doudna, 2015). The continuing application of Cas9 technologies to various studies has stimulated strong interest in the molecular basis by which Cas9 recognizes and cleaves its target DNA.

A series of crystal structures of Cas9 with and without the sgRNA and the target DNA have been solved (Anders *et al*, 2014; Jinek *et al*, 2014; Nishimasu *et al*, 2014; Jiang *et al*, 2016, 2015). These structural studies demonstrated the multi-domain architecture of Cas9, which mainly consists of a recognition (REC) lobe and a nuclease (NUC) lobe. The NUC lobe can be further divided into the HNH, RuvC and PAM-interacting (PI) domains. The crystal structures also revealed the sequential rearrangements of the Cas9 domains upon binding to the sgRNA and the target DNA. The binding of the sgRNA induces a large rotation of the REC lobe to convert Cas9 into the active conformation to form a central channel, which can accommodate the sgRNA-target DNA heteroduplex. Along with the DNA binding, the PI domain recognizes the PAM sequence in the target DNA, leading to the heteroduplex formation (Anders *et al*, 2014). This heteroduplex formation induces the translocation of the HNH domain and conformational changes in the RuvC domain, to cleave the double strands of the target DNA. These domain rearrangements during the Cas9 catalytic processes have been further confirmed by bulk FRET measurements (Sternberg *et al*, 2015).

Although the structural studies have revealed the distinct Cas9 domain configurations of the apo, sgRNA-bound and sgRNA/DNA-bound states, the crystal structures have also shown that some parts of Cas9 are disordered, suggesting that the domain configurations are flexible under specific conditions (Nishimasu *et al*, 2014; Jiang *et al*, 2016). The crystal structures and bulk FRET measurements indicated that the position of the HNH domain in the sgRNA/DNA-Cas9 ternary complex flexibly translocates relative to the REC lobe (Nishimasu *et al*, 2014; Anders *et al*, 2014; Sternberg *et al*, 2015; Jiang *et al*, 2016). In all of the available crystal structures, the active site in the HNH domain is located away from the cleavage site of the target DNA (Nishimasu *et al*, 2014; Anders *et al*, 2014; Jiang *et al*, 2016). Thus, the transition of the HNH domain that leads Cas9 to adopt conformations beyond those solved by the crystal structures should be crucial for the DNA cleavage. In addition, mismatched base pairs in the sgRNA-DNA heteroduplex hamper the HNH transition (Sternberg *et al*, 2015; Dagdas *et al,* 2017), suggesting that the flexibility of the HNH domain is closely related to not only DNA cleavage but also DNA recognition. A previous single molecule study implied the conformational flexibility during the DNA binding process (Singh *et al*, 2016), and molecular dynamics simulations have also shed light on the importance of the flexible movements of the Cas9 domains in the sgRNA/DNA binding (Zuo and Liu 2016; Palermo *et al.* 2016; Zheng, 2017). Thus, the flexibility of the Cas9 domain configuration could be an important factor in the Cas9 catalytic processes. However, direct experimental evidence of such flexible movement of the Cas9 domain in solution has not been reported.

To address this question, we directly observed the movement between the REC-RuvC, REC-HNH and HNH-RuvC domains, using single-molecule FRET (smFRET). Even in the steady state in the presence or absence of nucleic acids, a subset of Cas9 molecules demonstrated dynamic fluctuations in the FRET efficiency, providing strong evidence that the Cas9 domains move in a flexible and reversible manner. Further analysis suggested that the HNH domain accesses the DNA cleavage site only during the flexible domain movements, yielding new insights into the molecular basis of the Cas9 catalytic process.

## RESULTS

### Experimental setup for single-molecule FRET measurements of Cas9

To directly observe the mobility of the Cas9 domains at the single molecule level, Cas9 was site-specifically labeled with Cy3 and Cy5 fluorochromes. Using C80L/C574E cysteine-free Cas9, which has activity comparable to wild-type Cas9 (Nishimasu *et al*, 2014), as the starting construct, we introduced three pairs of cysteine residues at D435/E945, S355/S867 and S867/N1054 in Cas9, as done in a previous bulk FRET study (Sternberg *et al*, 2015). These three FRET constructs were designed to monitor the movements between REC-RuvC (D435C-E945C), REC-HNH (S355C-S867C) and HNH-RuvC (867C-N1054C), respectively (Fig 1A-C). The introduced cysteine residues were labeled with Cy3-(donor) and Cy5-(acceptor) maleimide. Furthermore, the constructs were genetically fused with biotin-carboxyl-carrier-protein (BCCP) at their N-terminus, to anchor the Cas9 molecules on a glass surface via an avidin-biotin linkage (Fig 1E). We first examined whether the FRET constructs retain their catalytic activity. All three BCCP-tagged fluorescent Cas9 constructs showed over 90% DNA cleavage activity as compared with wild-type Cas9 (Fig EV1), confirming that the cleavage activity is not substantially affected by the fluorescent labeling and the fusion with the BCCP-tag.

**Figure 1.**
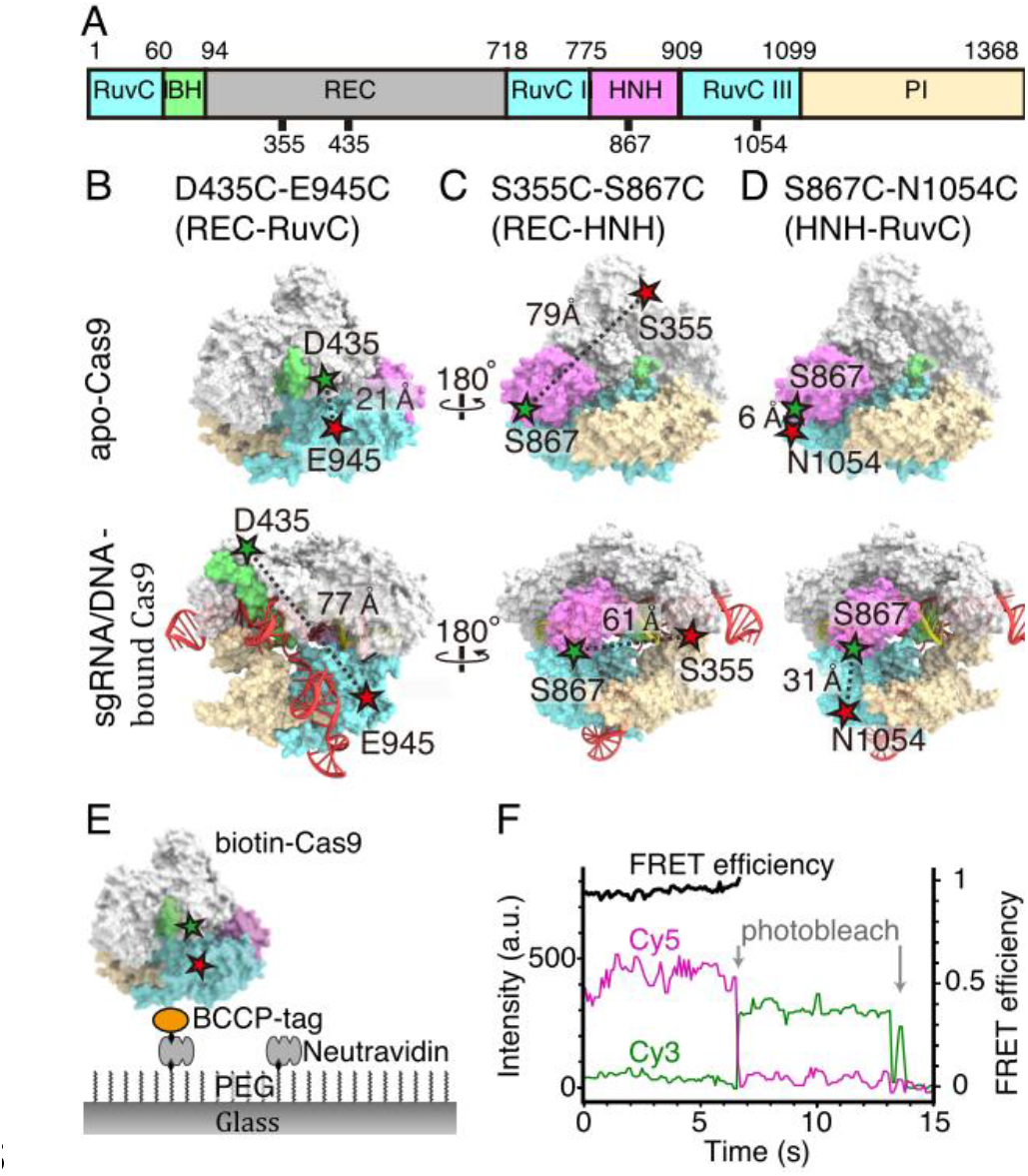
Experimental setup for smFRET measurement of Cas9 domain smovements. A The sequence diagram of the Cas9 molecule. The numbers indicate the amino acids that were fluorescently labeled in this study. B-D Designs of Cas9 for single molecule FRET (smFRET) measurements. We constructed three constructs: D435C-E945C (B), S355C-S867C (C) and S867C-N1054C (D). Surface rendered models of Cas9 were generated from PDB 4CMP for apo-Cas9 (upper models) and PDB 4OO8 for sgRNA/DNA-bound Cas9 (bottom models). HNH-domain, REC lobe, RuvC domain, PI domain and Bridge helix are colored pink, gray, blue, light brown and green, respectively. The Cy3-and Cy5-labeled amino acids are depicted by green and red stars. E Schematic drawing of the smFRET measurement system. Cas9, biotinylated via BCCP (Biotin Carboxyl Carrier Protein), was immobilized on a PEG (polyethylene glycol)-and biotin-PEG-coated glass surface, using the avidin-biotin system. Images are not to scale. F Time trajectories of single-molecule FRET efficiency of the D435C-E945C construct, labeled with Cy3 and Cy5. The green and magenta lines represent the fluorescence intensities of Cy3 and Cy5, respectively. We calculated the FRET efficiency (black lines) from the intensities of Cy3 and Cy5 before the photobleaching of either fluorochrome.

We then performed smFRET measurements of the fluorescently labeled Cas9 molecules under nucleic-acid free, sgRNA-bound and sgRNA/DNA-bound conditions, using total internal reflection fluorescent microscopy (TIRFM). To ensure the binding states of the Cas9 molecules in each condition, we incubated 0.3 to 1 nM fluorescently labeled Cas9 and 200 nM sgRNA with or without 200 nM target DNA, to measure the sgRNA-bound and sgRNA/DNA-bound Cas9 molecules. Considering the saturation rate of sgRNA on Cas9 (Fig EV2) and the dissociation constant value (*Kd*) of 0.8 nM for the target DNA loading onto sgRNA-bound Cas9 (Sternberg *et al*, 2015), we assumed that almost all of the fluorescently labeled Cas9 molecules were occupied with nucleic acids under our assay conditions. The sgRNA/DNA-bound molecules in our assay should maintain the ternary complex of the sgRNA and the cleaved target DNA, because previous studies demonstrated that Cas9 cleaves the target DNA at a rate higher than 10 min^-1^ and remains tightly bound to the cleaved DNA (Jinek *et al*, 2012; Sternberg *et al*, 2014; Sternberg *et al*, 2015). The Cas9 molecules were then anchored on the glass surface through BCCP, and illuminated with a 532-nm laser under TIRFM. The FRET efficiency of each Cas9 molecule was calculated from the recorded fluorescence intensities of Cy3 and Cy5 (Fig 1F). After the smFRET measurements, we confirmed that 68–95% of the observed Cas9 molecules labeled with Cy3 and Cy5 showed FRET under the tested conditions (Table EV1), using the acceptor bleaching method (see Method Details). Thus, we further analyzed the FRET trajectories of Cas9 molecules that showed FRET.

### Dynamic rearrangements of the Cas9 domains upon nucleic-acids binding

From the FRET efficiency of the Cas9 molecules (Fig 2), we validated the dynamic rearrangements of the Cas9 domains upon sgRNA and target DNA binding. In the apo state (Fig 2A), the FRET histograms of the fluorescently-labeled D435C-E945C (left panel) and S867C-N1054C (right panel) showed primary peaks at 0.99 ± 0.02 and 0.99 ± 0.06 (median ± HWHM) FRET efficiencies, respectively. Consistent with the crystal structures (Fig 1A, C and Table EV1), the high FRET efficiencies of the constructs indicated the close locations between the labeled amino acids. In contrast, the FRET histogram of the S355C-S867C construct in the apo state showed the primary peak at 0.12 ± 0.08 (Fig 2A, center panel), indicating the longer distance between the labeled amino acids. Upon sgRNA binding, the FRET efficiency of the D435C-E945C construct decreased (Fig 2B, left panel), suggesting that sgRNA binding induced drastic rotation of the REC lobe relative to the RuvC domain. In contrast, the changes in the FRET efficiencies of S355C-S867C and S867C-N1054C for the sgRNA binding were only slight (Fig 2B), as estimated from the crystal structures (Table EV1). Subsequent DNA binding increased the FRET peak values of D435C-E945C to 0.25 ± 0.11 and 0.98 ± 0.04 (Fig 2C, left panel). Similarly, the S355C-S867C construct exhibited an increase in the FRET efficiency upon the target DNA binding (Fig 2B, center panel), suggesting that the HNH domain approaches the cleavage site of the target DNA. This model of the HNH domain transition was further supported by the appearance of a low FRET distribution (0–0.5 FRET efficiency) in the histogram of the S867C-N1054C construct with the sgRNA and the target DNA (Fig 2C, right panel). Note that the changes in both the distance and orientation between the domains would contribute to the FRET efficiency shifts, because the fluorochromes on the Cas9 molecules showed high anisotropy (0.34-0.41 for Cy3 and 0.27-0.32 for Cy5, Appendix Fig S1). However, the timing and direction of the shifts were consistent with the previously proposed model (Nishimasu *et al*, 2014; Jinek *et al*, 2014; Jiang *et al*, 2015, 2016).

**Figure 2.**
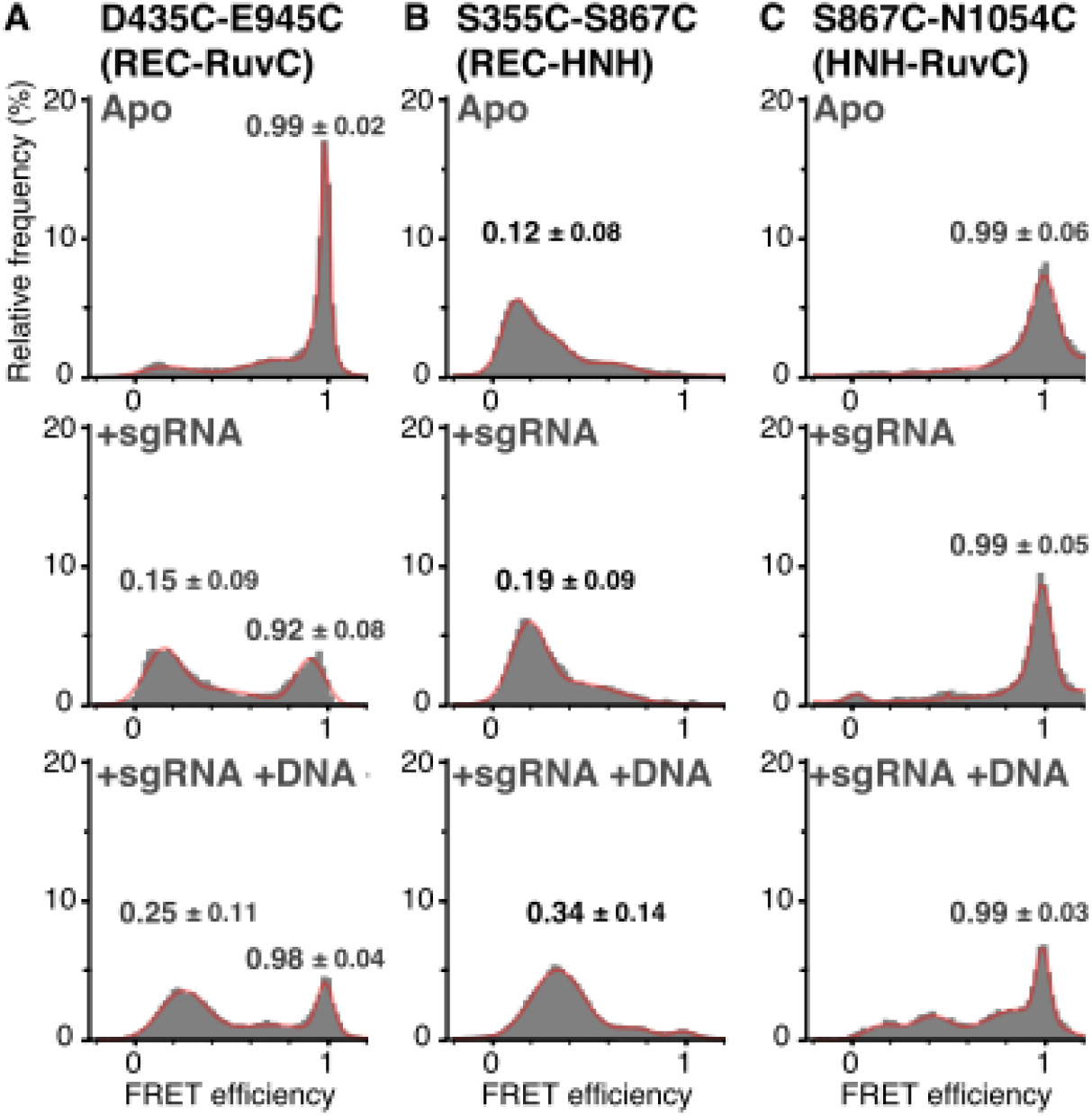
FRET efficiency histograms of all measured Cas9 molecules. A-C FRET efficiency histograms of the D435C-E945C (left panels), S355C-S867C (center panels) and S867C-N1054C (right panels) constructs. The histograms were generated from the time traces of the FRET efficiency in the absence of nucleotides (A), in the presence of 200 nM sgRNA (B) and in the presence of 200 nM sgRNA and 200 nM plasmid DNA (C). All of the experiments shown in this figure were performed in the presence of Mg^2+^. The numbers of measured molecules are summarized in Table EV1. The histograms were fitted with multi-peaks Gaussian curves (red). The peak values of the primary peaks of FRET efficiency are shown on the histograms (median ± HWHM).

### The Cas9 domains showed highly flexible and reversible movements

The histograms of the FRET efficiency under all of the tested conditions did not exhibit simple single-peak distributions (Fig 2), suggesting that the distances and/or angles between the Cas9 domains are not fixed. Consistently, a fraction of Cas9 molecules showed frequent fluctuations in the FRET efficiency between multiple FRET states (Fig 3). These fluctuations indicate the highly flexible and reversible movements of the Cas9 domains and represent the direct observation of the Cas9 domain fluctuations in solution.

**Figure 3.**
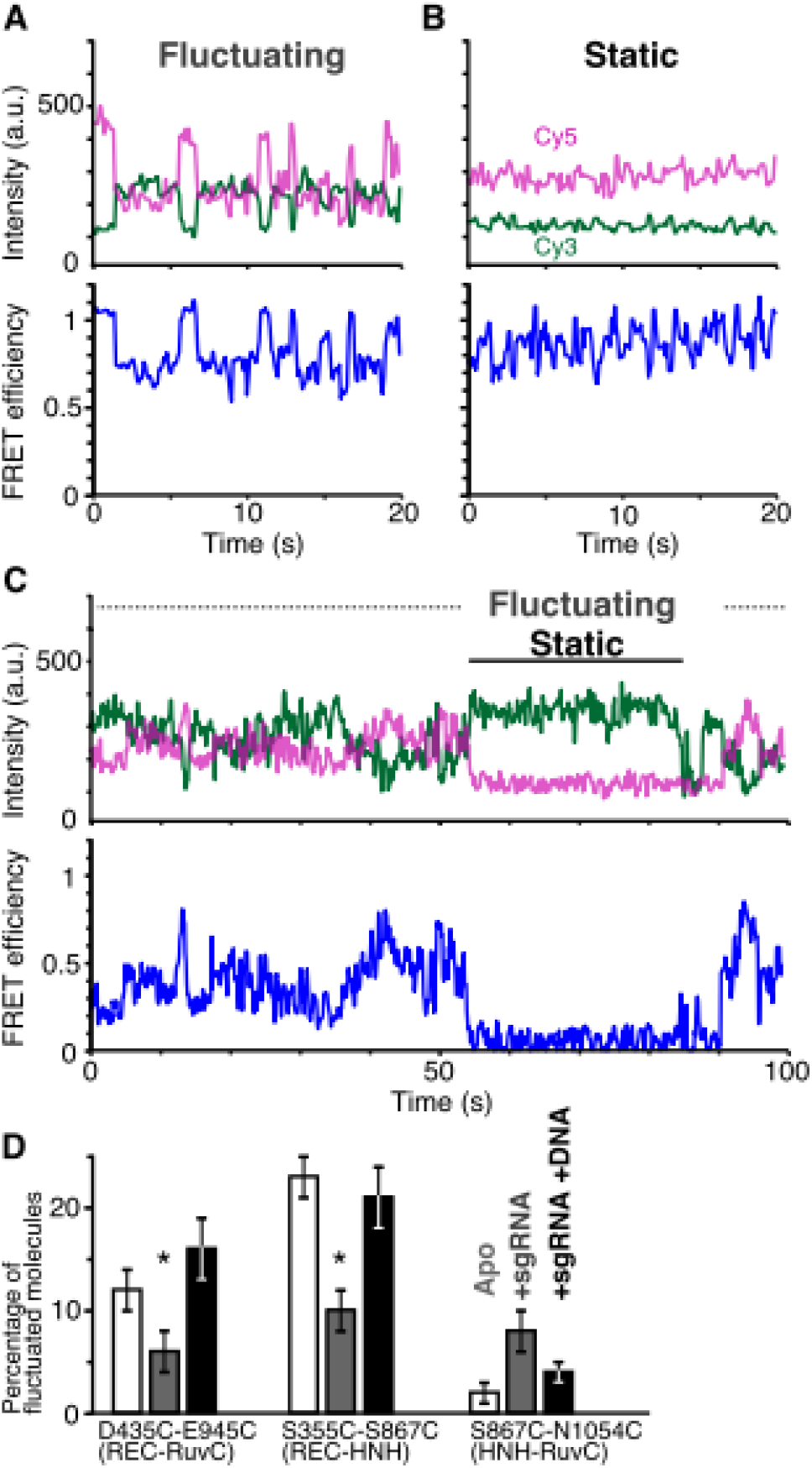
The binding of sgRNA and target DNA changes the flexibility of the Cas9 domains. A, B Representative time trajectories of fluctuating (A) and static (B) D435C-E945C molecules in the sgRNA/DNA-bound ternary complex labeled with Cy3 and Cy5. The green and magenta lines represent the fluorescence intensities of Cy3 and Cy5, respectively (top trace). We calculated the FRET efficiency (black lines) from the intensities of Cy3 and Cy5 (bottom trace). C Some of the time trajectories of the fluorescence intensities (top trace) and the single-molecule FRET efficiency (bottom trace) show both fluctuating of and static phases. D The percentage of Cas9 molecules that showed fluctuations in FRET efficiency. The numbers of measured molecules are summarized in Table EV1. The bars from left to right represent the percentages in the absence of nucleic acid (white), in the presence of 200 nM sgRNA (grey), and in the presence of 200 nM sgRNA and 200 nM plasmid DNA (black). Error bars show SEM. Asterisks indicate the statistical differences (*P* < 0.05, Steel-Dwass test).

During the 100-s observations, some molecules exhibited transitions between the static and fluctuating states (Fig 3C), suggesting that the Cas9 domains are in equilibrium between these states. Since the flexibility of the Cas9 domains should affect this equilibrium, we considered the percentage of fluctuating molecules to be an indicator of the domain flexibility (Fig 3D). Here, we defined a fluctuating molecule as a fluorescently-labeled Cas9 that showed more than two anti-correlated shifts in the fluorescence intensities of Cy3 and Cy5 during our observation period (see Materials and Methods).

The percentage of fluctuating molecules depended on the binding state of Cas9 (Fig 3D). As a common property of the D435C-E945C and S355C-S867C constructs, sgRNA binding lowered the percentage, suggesting the decreased flexibility between the REC and NUC (the HNH and RuvC domains) lobes by sgRNA binding. In contrast to the sgRNA binding, the target DNA binding increased the percentage of fluctuating molecules for both constructs (Fig 3D), suggesting the increased flexibility between the REC and NUC lobes in the Cas9-sgRNA-DNA ternary complex. Although the FRET fluctuations could be brought about by the increased dynamics within the REC domain itself, because the two opposite positions in the REC domain (S435 and S355) showed similar tendencies in their flexibility, it is most likely that the flexible movements occur between the two lobes. We further analyzed the flexibility in the NUC lobe, using the S867C-N1054C construct. Unlike the flexibility between the REC and NUC lobes, the flexibility between the HNH and RuvC domains apparently increased upon the sgRNA binding, but the differences were not statistically significant (*P* = 0.08, Steel-Dwass test). As compared with the D435C-E945C and S355C-S867C constructs, S867C-N1054C showed a relatively low number of fluctuating molecules (Fig 3D); however, there is a possibility that we underestimated the number of fluctuating molecules because, due to the short distance between S867 and N1054 (Nishimasu *et al*, 2014), the construct requires a relatively larger domain displacement for the FRET efficiency shift. Thus, it is not appropriate to compare the flexibilities of these three domains observed in the three constructs. However, because the percentages of fluctuating molecules of the D435C-E945C and S355C-S867C constructs were highly dependent on the nucleic-acid binding state, we conclude that the binding of nucleic-acids regulates the flexibility, at least between the REC and NUC lobes.

To elucidate the conformational differences between the fluctuating and static Cas9 molecules, we compared their FRET histograms (Fig 4 and EV3). We found that the FRET efficiency of fluctuating D435C-E945C molecules in the apo state was widely distributed from 0 to 1, in contrast to the very narrow FRET distribution (HWHM = 0.02) of static molecules in the apo state (Fig EV3). Considering the appearance of the low FRET peak in the FRET distribution of sgRNA-bound D435C-E945C (Fig 2B), some of the fluctuating molecules in the apo state should adopt a conformation that resembles the sgRNA-bound active form of Cas9. A similar tendency was observed in the S355C-S867C and S867C-N1054C constructs. The fluctuating S355C-S867C molecules in the apo state showed widely distributed FRET efficiencies without clear Gaussian peaks (Fig 4). In contrast, the static molecules showed a narrow peak at ~0.2 FRET efficiency (mean ± HWHM = 0.17 ± 0.07) in the apo state, and a gradual increase of the efficiency by the sgRNA binding. In the case of the S867C-N1054C construct, the FRET distribution of the static molecules showed a narrow peak at a high FRET efficiency (mean ± HWHM = 0.99 ± 0.06) in the apo state, and the gradual appearance of a low FRET population upon sgRNA and target DNA binding (Fig EV3). In contrast, the fluctuating S867C-N1054C molecules frequently showed low FRET efficiencies in the apo and sgRNA-bound states. These results demonstrate that flexible domain movements allow Cas9 to adopt different conformations from the static ones solved in the crystal structures.

**Figure 4.**
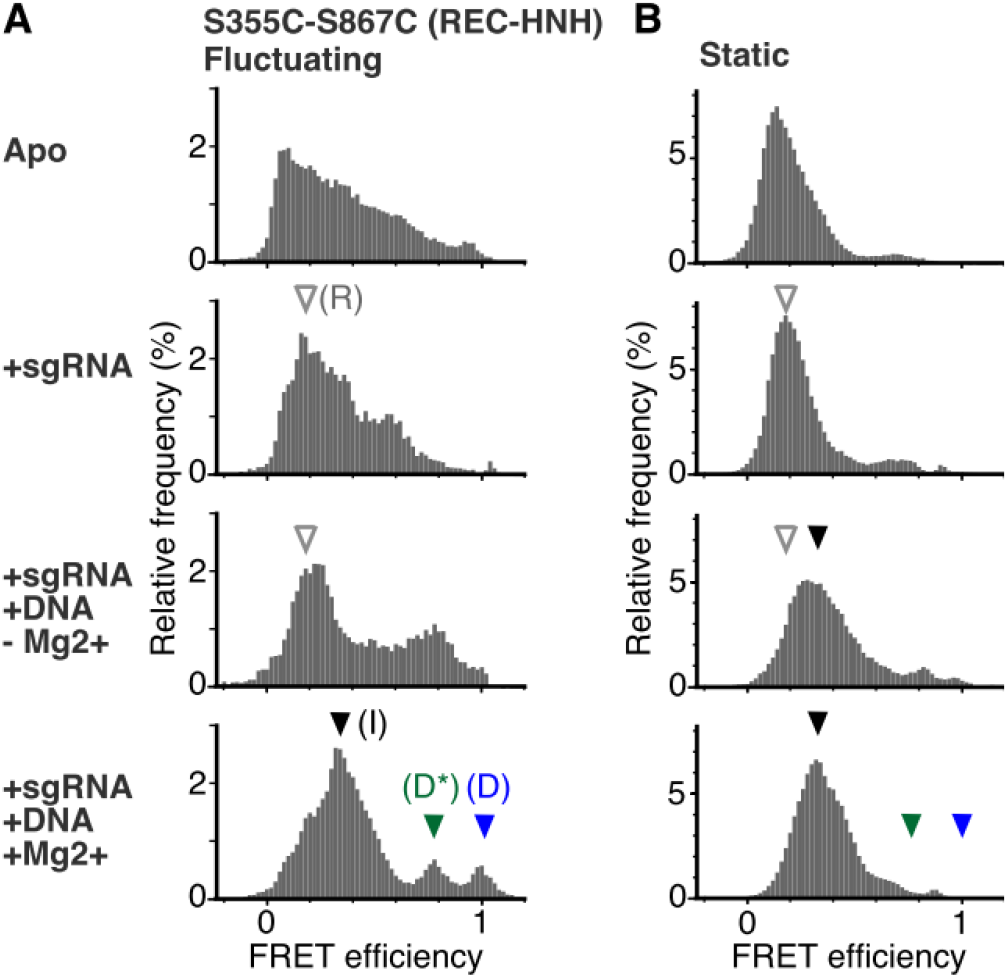
The HNH domain transiently exists at the cleavage competent D position during the flexible movement. A, B FRET efficiency histograms of fluctuating (A) and static (B) S355C-S867C molecules. The histograms were generated from the time traces of the FRET efficiency. The numbers of measured molecules are summarized in Table EV1. The panels from top to bottom show data in the absence of nucleic acids, in the presence of 200 nM sgRNA, in the presence of 200 nM sgRNA and 200 nM plasmid DNA without Mg^2+^, and in the presence of 200 nM sgRNA and 200 nM plasmid DNA with Mg^2+^, respectively. In the presence of Mg^2+^, the fluctuating molecules exhibited three clear FRET efficiency peaks, corresponding to the three HNH positions relative to the REC lobe (A). According to the structural and functional data, we refer to the HNH positions corresponding to the ~0.2 (white arrowhead), ~0.4 (black arrowhead), ~0.8 (green arrowhead) and ~1.0 (blue arrowhead) FRET efficiencies as the R, I, D* and D positions, respectively. The same FRET efficiencies are indicated by the arrowheads in the histograms of static molecules (B).

### The HNH domain accessed the DNA-cleavage position only during the flexible movement

To assess the effects of the flexible domain movements on the DNA cleavage process, we further analyzed the movements of the HNH domain in the Cas9-sgRNA-DNA ternary complex. The FRET efficiency distributions of the fluctuating S355C-S867C molecules in the ternary complex exhibited several clear peaks, in contrast to the widespread distributions of the apo-Cas9 and sgRNA-bound binary complex (Fig 4). These results suggest that the HNH domain in the ternary complex moves between distinct positions relative to the REC lobe. Consistently, previous studies have demonstrated that the ternary complex can adopt at least two conformations in which the HNH domain is close to or far from the cleavage site of the target strand (Nishimasu *et al*, 2014; Anders *et al*, 2014; Jiang *et al*, 2016).

Since Cas9 requires Mg^2+^ for DNA cleavage (Jinek *et al*, 2012), the Cas9-sgRNA-DNA complex can be trapped in the pre-cleavage state in the absence of Mg^2+^. The peak values of the FRET efficiency were ~0.2 and ~0.8 in the absence of Mg^2+^ (Fig 4A). As the lower peak value (~0.2) was similar to that of the sgRNA-bound S355C-S867C binary complex (Fig 2B), we considered the molecules with lower FRET efficiency as representing the RNA-bound (R) conformations, in which the HNH domain is located far from the target DNA (R position; distance between S355 and S867 ~8 nm, Table EV1). The higher FRET efficiency (~0.8) indicates that the HNH domain exists very close to the cleavage site, but the Cas9 molecules in the absence of Mg^2+^ do not cleave the target DNA. Therefore, we refer to the HNH position with the higher FRET efficiency as the DNA semi-docked pre-cleavage (D*) position. The time trajectories of the FRET efficiency suggested that the HNH domain in the ternary complex fluctuates among the R and D* positions in the absence of Mg^2+^.

The addition of Mg^2+^ to the ternary complex clearly changed the manner of the HNH fluctuations (Fig 4A). The addition of Mg^2+^ increased the percentage of fluctuating molecules more than two-fold (7 ± 1% to 20 ± 2%, mean ± SEM), and had only a slight effect on the FRET histogram of the S355C-S867C molecules remaining in the static state (Fig 4B). In contrast, fluctuating sgRNA/DNA-bound S355C-S867C molecules showed three major FRET efficiency peaks in the presence of Mg^2+^ (approximately 0.4, 0.8 and 1.0; Fig 4A). The addition of Mg^2+^ increased the primary peak value to ~0.4. This increase is consistent with the previous bulk FRET study (Sternberg *et al*, 2015). As the value of ~0.4 is in between the FRET peaks in the absence of Mg^2+^ (~0.2 and ~0.8), in the majority of Cas9 molecules, the HNH domain would be located at an intermediate (I) position between the R and D* positions. The probability of the HNH domain existing in the D* position decreased by the addition of Mg^2+^. Instead of the decrease of the ~0.8 FRET peak, the peak of the highest FRET efficiency (~1.0) appeared in the presence of Mg^2+^. The highest FRET efficiency was not observed in the absence of Mg^2+^, suggesting that the HNH domain can visit the third position only in the presence of Mg^2+^. Consistently, the probability of the HNH domain existing in the third position increased when the Mg^2+^ concentration was increased (Fig EV4). Increases in the Mg^2+^ concentrations also enhanced the DNA cleavage rate, yielding a strong correlation between the cleavage rate and the percentages of the Cas9 showing the highest FRET efficiency. These results indicate that the third position represents the conformation in which the HNH domain cleaves the target DNA. Thus, we refer to this HNH position as the DNA-docked cleavage competent (D) position. Importantly, very few Cas9 molecules in the static state showed the FRET efficiency corresponding to the D position (Fig 4B), suggesting that the flexible movement is critical for the HNH domain to be located at the cleavage-competent D position. The time trajectories of the FRET efficiency demonstrated frequent and reversible transitions between these three FRET states (Fig 5A), suggesting that the HNH domain fluctuates between the I, D* and D positions.

**Figure 5.**
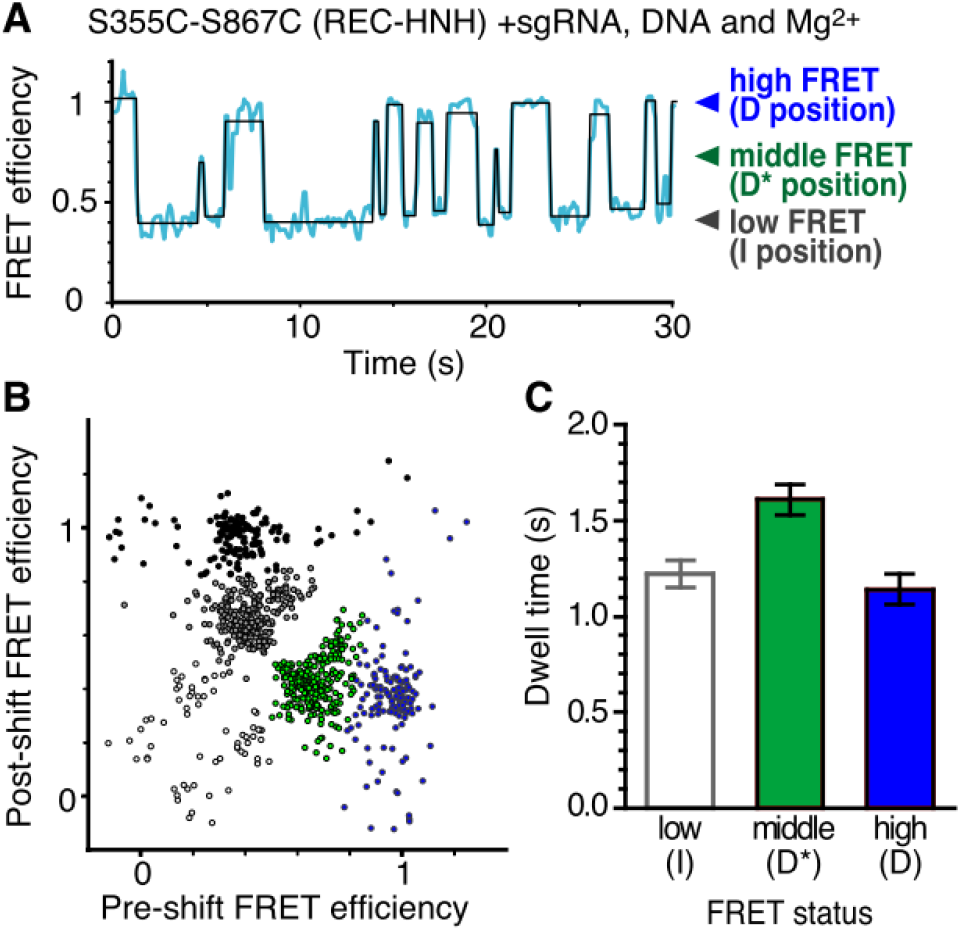
Reversible transitions of the HNH positioning in the ternary complex. A Representative time trajectory of the FRET efficiency, showing the fluctuation of the HNH domain in the sgRNA/target DNA-bound S355C-S867C complex with Mg^2+^. The transition points of the FRET efficiency (blue line) were detected using the HMM algorithm (black line). B The transition density plot of different FRET states of the sgRNA/target DNA-bound S355C-S867C complex with Mg^2+^. The density map was clustered into five groups (white, gray, black, green and blue closed circles) based on the *k*-means method with *k* = 5, suggesting that the HNH movement between the D* and D processes (middle and high FRET efficiencies) is rare. C Bar plot of the dwell times for each transition. The mean dwell times were determined by fitting the dwell time distributions (*n* = 399, 223 and 136 for low, middle and high FRET status, respectively) to a single exponential decay function (Appendix Fig S2). Error bars show SEM.

Finally, we investigated the movements of the HNH domain among the three positions. To analyze the relationship between the positions before and after the transitions of the HNH domain in the ternary complex, we measured the FRET time trajectories of sgRNA/DNA-bound S355C-S867C, using a hidden Markov model-based algorithm (Fig 5A), and plotted the FRET efficiencies of the pre- and post-HNH transitions (Fig 5B). Together with the transition density plot and Silhouette analysis (Fig EV5), the transitions can be classified into five types: transitions from a low FRET state to another low FRET state (I-R), between low and middle FRET states in both directions (I-D*) and between low and high FRET states in both directions (I-D). To our surprise, transitions between middle and high FRET states were rare (less than 2% of all transitions), suggesting that the HNH domain rarely moves between the D* and D positions, and therefore needs to adopt the undocked I position before relocating to the D* or D position.

Among the three positions, the HNH domain in the pre-cleavage D* position showed the longest dwell time before the transition (Fig 5C and Appendix Fig S2), suggesting the high stability of the HNH domain in the D* position, as compared to those in the other positions. Consistently, the frequency of the I to D* transition (219 times / 343 transitions = 64%) was approximately twice as high as that of the I to D transition (124 times / 343 transitions = 36%). Thus, the HNH in the D* position should be a thermodynamically stable conformation. However, as mentioned above, the HNH domain in the D position rarely moves to the D* position (Fig 5B). The results suggest that a structural barrier for the HNH transition exists between the D* and D positions, which must be collapsed by the transition to the I position.

## DISCUSSION

The purpose of the present study is to investigate whether Cas9 has a flexible structure in solution, as predicted by previous studies (Nishimasu *et al*, 2014; Jinek *et al*, 2014, Anders *et al*, 2014; Jiang *et al*, 2015; Sternberg *et al*, 2015; Jiang *et al*, 2016; Singh *et al*, 2016; Zheng, 2017). Here, using the smFRET technique, we directly observed the dynamic fluctuations of the Cas9 domain. These fluctuations allow Cas9 to adopt different conformations besides those previously reported by crystal structure analyses (Nishimasu *et al*, 2014; Jinek *et al*, 2014; Anders *et al*, 2014; Jiang *et al*, 2015; Jiang *et al*, 2016). Our detailed analysis highlights the potential roles of the transient conformations regulated by the flexibility in both the DNA cleavage and sgRNA/DNA binding processes.

Here, we summarize the flexibility of the Cas9 domain configuration observed in the present study (Fig 6). Judging from the percentages of the fluctuated molecules (Fig 3D), the NUC lobe flexibly moved relative to the REC lobe in the apo-Cas9. The binding of the sgRNA stabilizes the fluctuations between the REC and NUC lobes, but the subsequent target DNA binding enhances the fluctuations (Fig 3D). Our smFRET data indicated that the HNH domain in the ternary complex fluctuated between three distinct positions in the presence of Mg^2+^: the I, D* and D positions (Fig 5).

**Figure 6.**
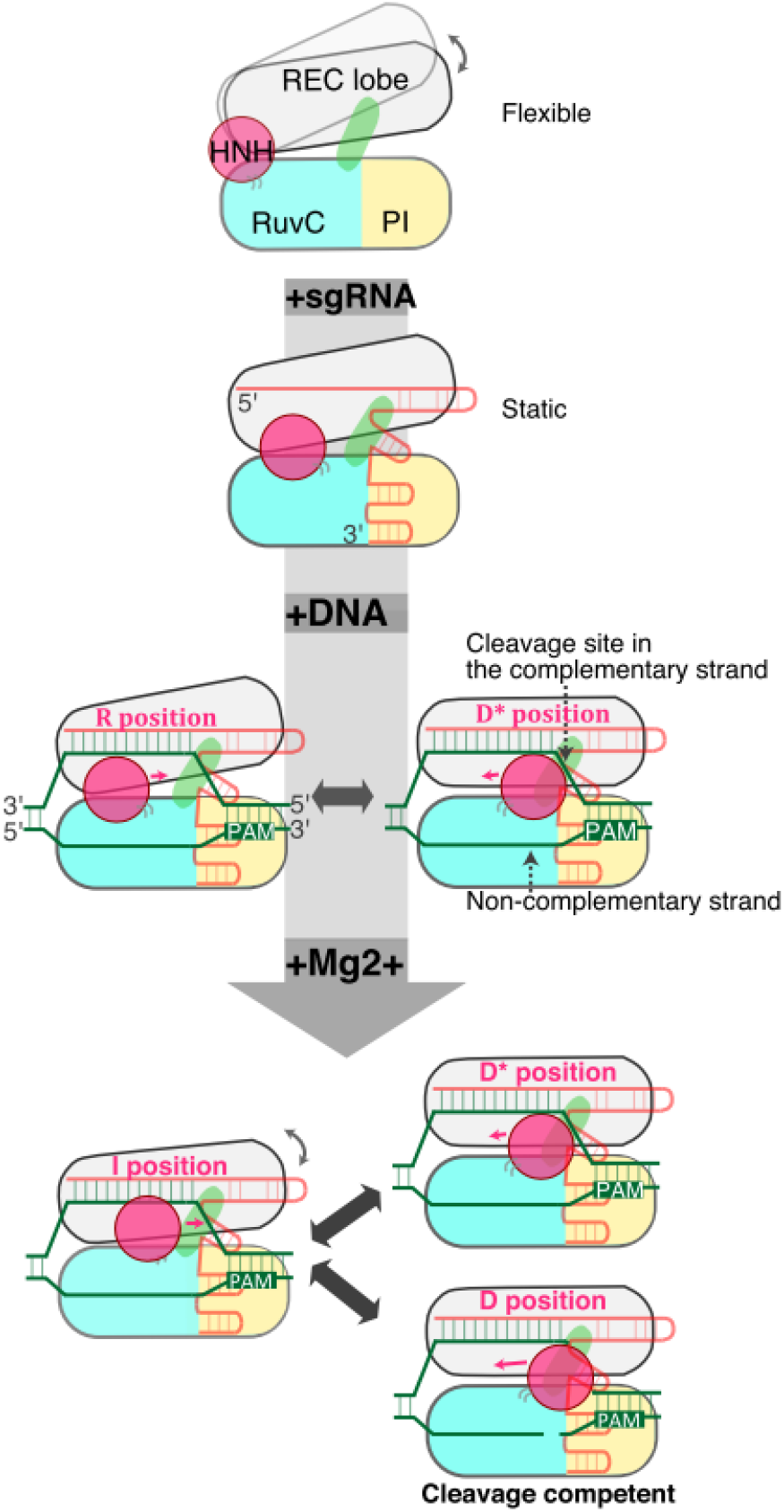
Model of Cas9-mediated DNA cleavage. The relative flexible movements of the REC lobe (gray) against the HNH (magenta) and RuvC (blue) domains are represented by the grey arrows. The binding of the sgRNA (orange) stabilizes the flexibility, but the binding of the target DNA (green) and Mg^2+^ increases the flexibility between the REC and NUC lobes. The HNH positions in the ternary complex are indicated by magenta letters.

Even in the presence of Mg^2+^, the Cas9 molecules in the static phase did not show the high FRET efficiency corresponding to the HNH domain in the cleavage competent D position (Fig 4B). This result indicated that the HNH domain can only access the D position during the fluctuating phase, thus emphasizing the importance of the flexible movement of the HNH domain in the DNA cleavage process. The movement of the HNH domain has been reported to control the nuclease activity of the RuvC domain on the noncomplementary strand, through intramolecular communication between the two domains (Sternberg *et al*, 2015; Jiang *et al*, 2016). Thus, besides its direct participation in the cleavage of the complementary strand, the flexibility of the HNH domain may also affect the noncomplementary strand cleavage by the RuvC domain.

The crystal structure demonstrated that apo-Cas9 adopts an autoinhibited conformation, in which the active sites in the HNH and RuvC domains are located away from the DNA binding cleft, and the interaction interfaces with the sgRNA are limited (Jinek *et al*, 2014). Our smFRET data revealed the fluctuations between the REC and NUC lobes in apo-Cas9 (Fig 3), indicating that apo-Cas9 adopts transient conformations in addition to the static conformations revealed by the crystal structure. The REC movement against the NUC lobe can provide additional interaction interfaces for the sgRNA; thus, the flexible movement in apo-Cas9 may play an important role in the assembly with the sgRNA.

After the submission of our manuscript, three preprints of similar smFRET studies have been posted on bioRxiv (Dagdas *et al*, 2017; Yang *et al*, 2017; Chen *et al*, 2017). Consistent with our data, these studies demonstrated the dynamic translocations of the HNH domain among multiple (R, I and D) positions, although the fluctuating and static molecules were not distinguished in these studies. There are also several discrepancies among the studies. For instance, the populations of S355C-S867C molecules showing the high FRET efficiency in the sgRNA/DNA-bound state are different among these studies. In the studies by Dagdas *et al* and Chen *et al*, most of the S355C-S867C molecules exhibited ~1.0 FRET efficiency, suggesting that almost all of the Cas9 molecules have the HNH domain in the cleavage competent (D) position. In contrast, the major peak value of the FRET efficiency was ~0.4 in the study by Yang *et al* and our study, suggesting that most of the HNH domain is located in the intermediate (I) position. The report by Yang *et al* and our study also demonstrated the existence of the pre-cleavage (D*) HNH position. Yang *et al* proposed the possibility that heparin, which was only included in the buffers used by Dagdas *et al* and Chen *et al*, produces the difference, but further analyses are required to understand the underlying cause of the discrepancy.

Although verification of the function of the flexible movements awaits further studies, our results open a new door toward modifying and expanding Cas9-based tools by modulating the domain flexibility. Since the HNH domain in the D* position must return to the I position before it translocates to the cleavage competent D position, mutations in the interface of the HNH domain and the REC lobe that destabilize the HNH domain in the D* and I positions may facilitate the HNH translocation to the D position, enhancing Cas9-mediated DNA cleavage. Together with the demonstration of the domain flexibility of apo-Cas9, which may play a role in the sgRNA binding, our data provide useful information for future improvements in Cas9-based tools for gene-editing, gene-visualization and gene expression control.

## MATERIALS AND METHODS

### Sample preparation

Since the C80L/C574E mutations in Cas9 do not affect the cleavage activity and improve the solution behavior (Nishimasu *et al*, 2014), we used the Cas9 C80L/C574E mutant as wild-type Cas9 in this study. We introduced the mutations into the Cas9 C80L/C574E mutant, to prepare D435C-E945C, S355C-S867C and S867C-N1054C. The Cas9 proteins were prepared as previously described (Nishimasu *et al*, 2014), with minor modifications. Briefly, the Cas9 variants were expressed as His6-GST-fusion proteins at 20°C in *Escherichia coli* Rosetta 2 (DE3) (Novagen), and purified by chromatography on Ni-NTA Super flow resin (QIAGEN). The His6-GST tag was removed by TEV protease digestion, and the proteins were further purified by chromatography on Ni-NTA, HiTrap SP HP (GE Healthcare), and Superdex 200 Increase (GE Healthcare) columns. The purified Cas9 was stored at −80°C until use.

### *In vitro* cleavage assay

*In vitro* cleavage experiments were performed as previously described (Anders *et al*, 2014), with minor modifications. A *Eco*RI-linearized pUC119 plasmid (100 ng, 5 nM), containing only one 20-nt target sequence followed by the NGG PAM (Appendix Fig S3), was incubated at 37°C for 5 min with the Cas9-sgRNA complex (25 and 50 nM) in 10 μL of reaction buffer, containing 20 mM HEPES-NaOH, pH 7.5, 100 mM KCl, 2 mM MgCl2, 1 mM DTT, and 5% glycerol. We confirmed that the plasmid DNA does not contain long off-target sequences (Appendix Fig S3). The reaction was stopped by the addition of a solution containing EDTA (40 mM final concentration) and Proteinase K (1 mg/mL). Reaction products were resolved on an ethidium bromide-stained 1% agarose gel and then visualized using an Amersham Imager 600 (GE Healthcare). For the cleavage assays at various Mg^2+^ concentrations, an *Eco*RI-linearized pUC119 plasmid (3.5 nM) was incubated at 25 °C for 30 min with the fluorescent Cas9 (S355C-S867C)–sgRNA complex (50 nM), in 10 μL of reaction buffer containing 20 mM HEPES-NaOH, pH 7.5, 100 mM KCl, 0.5 mM EDTA, 1 mM DTT, and 5% glycerol, with 0.5, 1, 2 and 5 mM MgCl2. Following electrophoresis on a 1.5 % agarose gel, the reaction products were fluorescently stained using Midori Green Advance (Nippon Genetics Co., Ltd.) and then visualized using a Typhoon FLA 9500 imager (GE Healthcare) equipped with a 473 nm excitation laser and an LPB filter (GE Healthcare).

### Preparation of the sgRNA and the target plasmid DNA

The sgRNA was transcribed *in vitro* with T7 RNA polymerase, using a PCR-amplified DNA template, and purified by 10% denaturing (7 M urea) PAGE. The target plasmid DNA was amplified in the *E. coli* DH5a strain, grown in LB medium (Nacalai Tesque, Inc., Japan) at 37°C overnight. The plasmid DNA was purified using a Midiprep kit (FastGene Xpress Plasmid PLUS Kit, NIPPON Genetics), according to the manufacturer’s method. The concentration of purified plasmid DNA was determined based on the absorption at 260 nm, using a NanoDrop 2000c spectrophotometer (Thermo Fisher). The single-guide RNA and the plasmid DNA were stored at -80 °C and -30 °C until use, respectively.

### Fluorescent labeling of Cas9

Cas9 was fluorescently labeled using Cy3-and Cy5-maleimide (GE Healthcare), according to the general method. Briefly, the buffer for Cas9 was exchanged into the labeling buffer (20 mM HEPES-KOH, pH 7.0, 100 mM KCl, 2 mM MgCl2, 5% glycerol), using a spin-gel filtration column (Micro Bio-Spin 30, Bio-Rad). Next, the Cas9 solution was incubated on ice for 30 min, after the final 0.5 mM TCEP addition into the Cas9 solution. Then, Cy3-and Cy5-maleimide were mixed with the Cas9 solution at a 1: 20 molar ratio between the protein and each dye. The maleimide labeling reaction was conducted on ice for 2 h. Excess fluorescent maleimide dye was removed twice, using assay buffer (AB: 20 mM HEPES-KOH, pH 7.5, 100 mM KCl, 2 mM MgCl2, 5% glycerol, 0.5 mM EDTA, 1 mM DTT) and spin-gel filtration columns (Micro Bio-Spin 30, Bio-Rad). The fluorescently labeled Cas9 was snap-frozen in liquid nitrogen and stored at -80 °C until use.

### FRET measurements for the stoichiometry of sgRNA binding to Cas9

All fluorescence measurements used a reaction mixture of 20 nM fluorescent Cas9 (D435C-E945C) with or without the sgRNA (10 nM, 20 nM, 50 nM, 100 nM or 200 nM) in AB with 0.1 U/μL RNasin Plus (Promega), and a commercial oxygen scavenger system (Pacific Bioscience) containing 2.5 mM TSY, 2.5 mM protocatechuic acid (PCA) and 50-fold diluted protocatechuic acid dehydrogenase (PCD) solution. Measurements were performed using a fluorescence spectrometer (RE-6000, Shimadzu, Japan) and a quartz cuvette with a 50 μL volume (T-703M-ES-10.50B, TOSOH, Japan), with 532 nm excitation and a scanning speed of 60 nm/min in the wavelength range of 550 nm to 750 nm in 1 nm increments, at room temperature.

### Perrin plot to determine the orientation factors

All fluorescence measurements using the reaction mixture of 100 nM fluorescent Cas9 (D435C-E945C, S355C-S867C and S867C-N1054C with no nucleic acid) in buffer (20 mM HEPES-KOH, pH 7.5, 100 mM KCl, 2 mM MgCl2, 0.5 mM EDTA, 1 mM DTT and a commercial oxygen scavenger system) with or without methyl cellulose (0, 0.001, 0.01 or 0.1%) were performed at room temperature, using the same fluorescence spectrometer and cuvette described in the previous section. The orientation factor *κ*^2^ was determined as described below, according to the previous method (Dale *et al*, 1979). Briefly, the fluorescence anisotropy measurement was performed by manually placing the polarization filters in front of the exciter and detector in the fluorescence spectrometer. For Cy3, the fluorescence intensity was measured at a wavelength of 566 nm by 554 nm excitation, while that of Cy5 was measured at a wavelength of 668 nm by 650 nm excitation. The slit width for emission and excitation was 5 nm, and the integration time was 1 s. Each measurement was repeated three times. Using these fluorescence intensities, the fluorescence anisotropy *r* was calculated as described below.

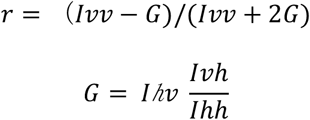

*Ivh* indicates the fluorescence intensity of the horizontal polarization excited by the vertical polarized light. *Ivv, Ihv* and *Ihh* are defined similarly. Following the plotting of 1/*r* against *T*/*η*, the y-intercept was calculated by fitting the plot to the linear function for each fluorescent Cas9, to estimate the *γ* values.

### Single molecule FRET measurement of fluorescent Cas9

The cover slips (No. 1S, 22 × 22 mm, Matsunami, Japan) were cleaned for 15 min, using 1 N KOH and an ultrasonic washing machine (BRANSONIC, Branson). All subsequent preparation procedures were performed on a clean bench (Matsusada Precision, Japan). After 20 rinses using Milli-Q water and drying in a dryer, the cover slips were cleaned using a plasma cleaner (YHS-R, SAKIGAKE-Semiconductor Co., Ltd., Japan or PR300, Yamato Scientific Co., Ltd., Japan). Next, the cover slips were completely dried in a dryer. Following the cleaning of cover slips as described above, one side of the cover slips was silanized by sandwiching 10 μL of N-2 (aminoethyl)-3-aminopropyl-triethoxysilane(KBE-603, Sin-Etsu Silicones, Japan. After an incubation at room temperature for 20 min, the cover slips were rinsed 20 times and dried. The silanized side of the cover slips was PEGylated by sandwiching 10 μL of 200 mg/mL NHS-PEG and 1 mg/mL NHS-PEG-biotin (BI-050TS, NOF, Japan) in 50 mM MOPS (pH 7.5) for the observed surface of a flow chamber, and 200 mg/mL NHS-PEG (ME-050-TS, NOF, Japan) in 50 mM MOPS (pH 7.5) was used for the non-observed surface (Yokota *et al.* 2009). Following an incubation at room temperature for 2 h under moist conditions, the cover slips were rinsed 20 times with Milli-Q water and completely dried. A 0.5 μL volume micro-chamber was made by placing a PEG-coated small coverslip of 11 mm × 11 mm, which was cut from a commercial coverslip, over a PEG-biotin coated 22 mm × 22 mm glass coverslip using double-sided adhesive tape (30 μm thickness, Nitto Denko, Japan) in a clean hood (Matsusada Precision Inc., Japan). First, 1 mg/mL Neutralized avidin (Wako, Japan) in AB was adsorbed onto the glass surface. After a 2 min incubation, the excess neutralized avidin was removed by 3 washes with 2 μL AB. Next, the glass surface in the micro-chamber was illuminated by a 532 nm laser for 40 s per one field using fluorescence microscopy, to photobleach any residual fluorescent particles on the glass surface. After 3 washes with 2 μL AB, 0.3 - 1 nM fluorescent Cas9 was adsorbed onto the glass surface, using the avidin-biotin interaction. Here, for the sgRNA-bound fluorescent Cas9 imaging, fluorescent Cas9 was incubated with a final concentration of 200 nM sgRNA for 2 min at room temperature in a 0.6 mL tube before the Cas9 absorption, while the fluorescent Cas9 was successively incubated with 200 nM sgRNA and 200 nM plasmid DNA for 2 min, for the sgRNA-and DNA-bound fluorescent Cas9 imaging. After a 2 min incubation and 3 washes with 2 μL AB, AB with a commercial oxygen scavenger system (Pacific Bioscience), containing 2.5 mM TSY, 2.5 mM protocatechuic acid (PCA) and 50-fold diluted protocatechuic acid dehydrogenase (PCD) solution was placed in the micro-chamber for all samples, and then 200 nM sgRNA was added for the sgRNA-bound fluorescent Cas9 imaging and 200 nM sgRNA and 200 nM plasmid DNA were added for the sgRNA- and DNA-bound Cas9 imaging. Finally, the micro-chamber was subjected to total internal reflection fluorescence microscopy (TIRFM) for the single molecule FRET (smFRET) measurements.

The smFRET measurements of fluorescent Cas9 were achieved using a Nikon Ti-E based TIRFM, equipped with a multi-band filter set for fluorescence microscopy (LF405/488/532/635-A, Semrock), a dual-view apparatus (Optical Insights) containing dichroic (630, Optical Insights) and emission filters (FF01-593/40-25 for Cy3 imaging and FF01-692/40-25 for Cy5 imaging, Semrock), and a back-illuminated EMCCD camera (Andor, iXon+). Illumination was provided by a 532 nm laser (Coherent, Sapphire) and a 642 nm laser (Coherent, Cube). Image acquisition for the smFRET measurements was performed with an acquisition rate of 10 frames per second, using 532 nm illumination and open source microscopy software (Micro-Manager, Open Imaging) (Edelstein *et al,* 2014). The FRET efficiency distributions were calculated over the duration of the photobleaching of the fluorescent dye (donor or acceptor) or the entire observation time (120 s for D435C-E945C and S355C-S867C; 40 s for S867C-N1054C), in cases where no photobleaching was observed. We typically collected data from 12 observation fields of at least three different chambers for each condition. Following the smFRET measurements, the same field was illuminated using a 642 nm laser to directly excite the Cy5 fluorescence, for counting the Cy3 and Cy5 double-labeled Cas9. This procedure allowed us to exclude the data from the molecules labeled with only the donor dye. From the Cy5 intensity before and after the photobleaching process, we judged whether the decreases in the Cy5 fluorescence intensity during the smFRET observation reflected the Cas9 conformational changes or were caused by fluorescence photobleaching. We confirmed that the levels of donor leakage in the acceptor detection channel and fluorescent photoblinking were negligible in our assay conditions.

For the smFRET analysis, the exported image data were imported into a home-built program written in Python and converted into fluorescence intensity, based on the fluorescent spots of both Cy3-and Cy5-labeled Cas9. The FRET efficiency of a single molecule was calculated as *I_A_*/(*I_A_*+γ*I_D_*) (Roy *et al*, 2008). Here, *I_A_* and *I_D_* are the fluorescence intensities of the acceptor and donor, respectively. *γ* is equivalent to |Δ*I_A_*/Δ*I_D_*|, where Δ*I_A_* and Δ*I_D_* are the fluorescence intensity changes of the acceptor and donor upon FRET efficiency fluctuation or photobleaching, respectively. The fluctuating molecules were initially sorted with standard deviation values of 0.4-Hz low pass filtered time traces. After the initial sort, traces with multiple FRET transitions within 2.5 s were re-categorized as the fluctuating state. The traces exhibiting both fluctuating and static states were categorized as the fluctuating molecule. The transition points in the fluctuating traces of the sgRNA/DNA-bound fluorescently labeled Cas9 (S355C-S867C) were detected based on the Hidden Markov Model (HMM) with the Baum-Welch forward-backward algorithm and the Viterbi algorithm (McKinney *et al*, 2006), using the hmmlearn library for Python (https://github.com/hmmlearn/hmmlearn). Here, we assumed that HMM has three states, according to the FRET efficiency distribution (the bottom histogram in Fig 4A). The transition density plot was visualized using a Python plotting library (Matplotlib; http://matplotlib.org), while the plotted density was clustered into five groups based on the k-means method with *k* = 5, using the machine learning package for Python (Scikit-learn; http://scikit-learn.org/).

### Data availability

The smFRET data were uploaded with the manuscript.

## ACKNOWLEDGEMENTS

This study is supported by JST, CREST (to S.U) and MEXT, Grants-in-Aid for Young Scientists (B), 15K18514 (to T.S.) and 17K15100 (to T.K.). We thank the members of the Uemura and Nureki laboratories for valuable discussions. We also thank M. Sugawa for technical assistance and P. Karagiannis for helpful discussions and comments on the manuscript.

## Author contributions

T.K., H.N., T.S., O.N. and S.U. designed the study. S.O. and T.K. collected and analyzed smFRET data; T.K. collected and analyzed bulk FRET data; K.I. and S.K. prepared the fluorescently-labeled protein; S.K. and T.K. performed functional analyses; and S.K., T.K., H.N., T.S., O.N. and S.U. wrote the paper. All authors discussed the results and commented on the manuscript.

## Conflict of interest

The authors declare no conflict of interest.

## Expanded View Figure Legends

**Figure EV1.**
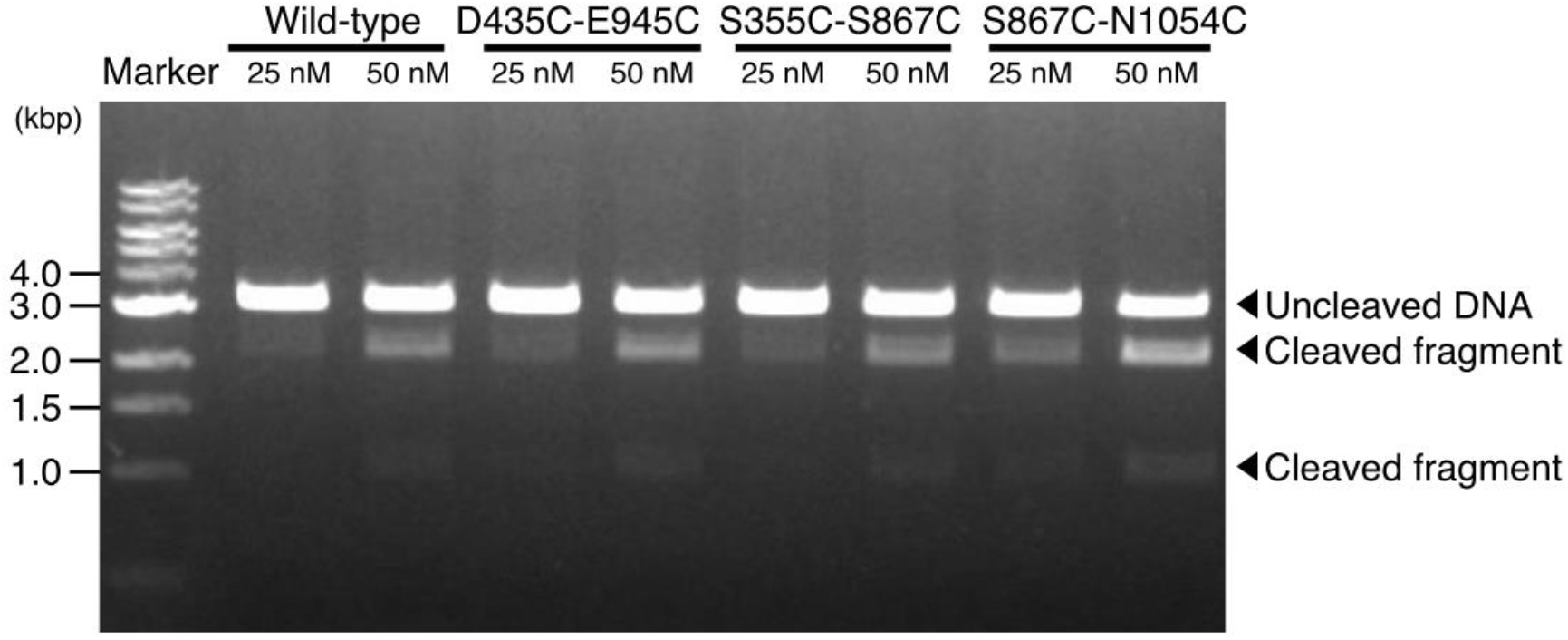
DNA cleavage activity of fluorescently-labeled biotin-Cas9. All three FRET constructs were labeled with Cy3 and Cy5 and were tested for nuclease activity. After an incubation of 25 or 50 nM Cas9 - sgRNA complex and 5 nM target DNA for 5 min at 37 °C, a fraction of the DNA was cleaved into two fragments. The three FRET constructs demonstrated nuclease activity comparable to that of non-labeled wild-type Cas9 (1.1 ± 0.1 for D435C-E945C, 0.9 ± 0.1 for S355C-S867C and 1.5 ± 0.3 for S867C-N1054C; mean relative activity ± SEM., n = 3).

**Figure EV2.**
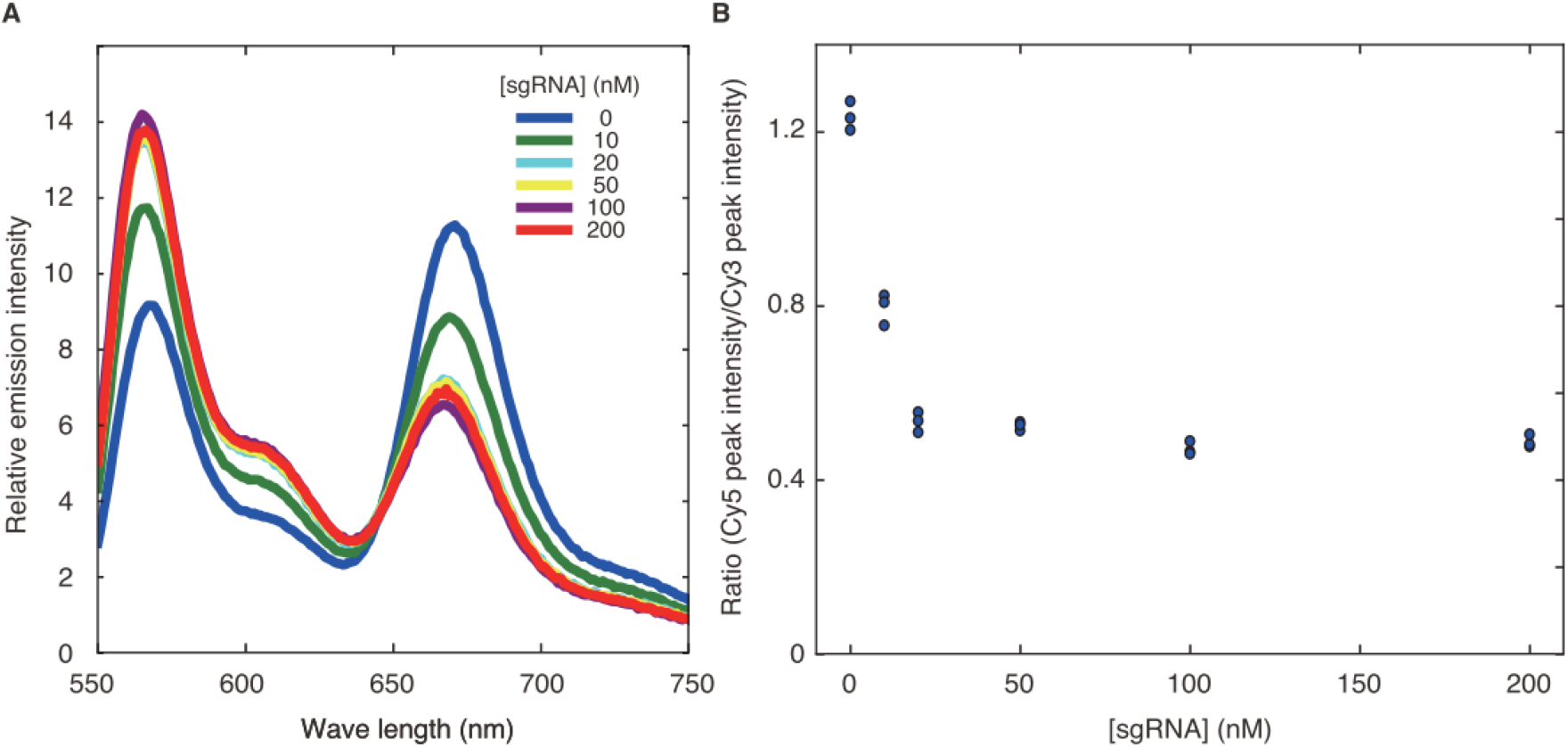
Stoichiometry of sgRNA binding to Cas9. A Fluorescence emission spectra of 20 nM fluorescent Cas9 (D435C-E945C) excited at 532 nm. The Cy3-and Cy5-fluorescence intensity changes were coupled with the FRET efficiency change, according to the sgRNA concentration. B Quantification of the ratio between Cy3-and Cy5-fluorescence intensities. The ratios of Cy5-fluorescence peak intensity over Cy3-fluorescence peak intensity were plotted against the sgRNA concentration (n=3 for each sgRNA concentration). The FRET efficiency change coupled with the sgRNA binding was almost saturated at the Cas9 to sgRNA ratio of 1:1.

**Figure EV3.**
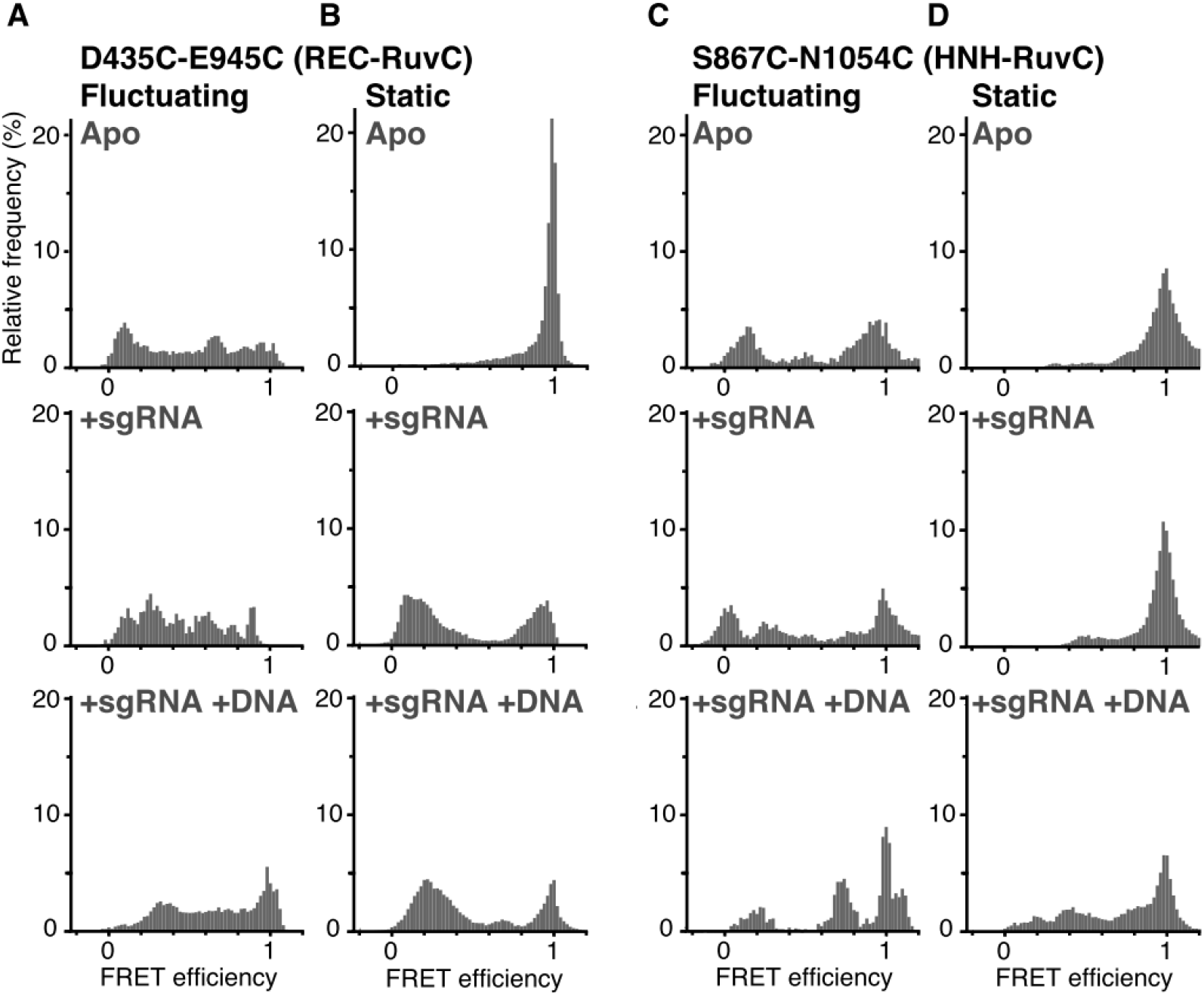
FRET efficiency histograms of fluctuating and static Cas9 molecules. A-D FRET efficiency histograms of fluctuating D435C-E945C (A), static D435C-E945C (B), fluctuating S867C-N1054C (C) and static S867C-N1054C (D). The numbers of measured molecules are summarized in Table EV1. The top, middle and bottom panels show data acquired in the absence of nucleotides, in the presence of 200 nM sgRNA and in the presence of 200 nM sgRNA and 200 nM plasmid DNA, respectively. All of the data in this figure were acquired in assay buffer containing 2 mM MgCl2.

**Figure EV4.**
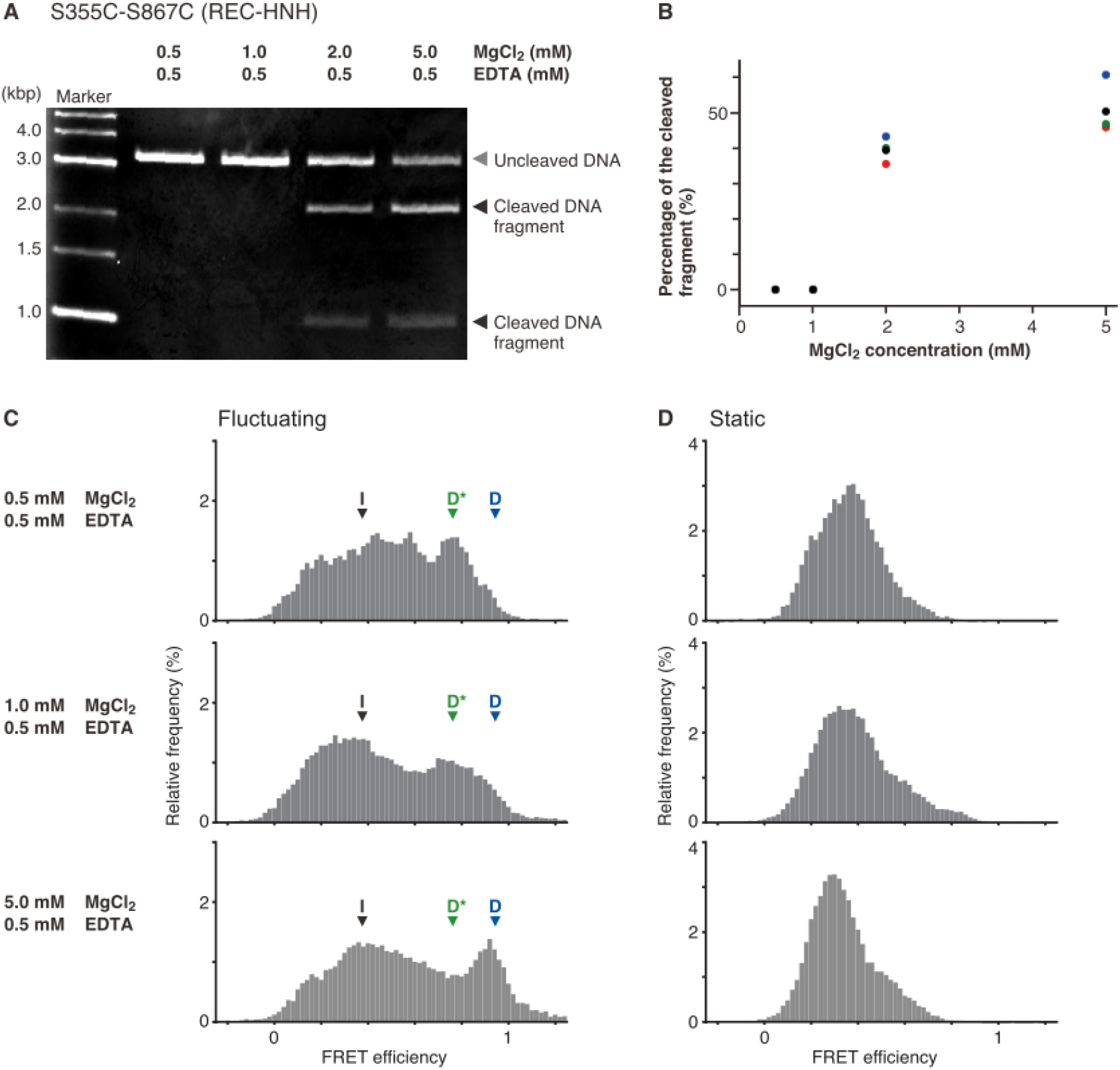
Effects of Mg^2+^ concentration on the DNA cleavage activity and the HNH location in the sgRNA/DNA-Cas9 ternary complex. A Representative gel image of the DNA cleavage assay using the fluorescently labeled S355C-S867C construct. The sgRNA/DNA-Cas9 ternary complex was incubated at room temperature (25 °C) for 30 min. This condition is equivalent to that of the smFRET measurement, as we observed smFRET for approximately 30-40 min at room temperature. B Percentages of cleaved DNA against MgCl2 concentration. The plot shows the results of four individual assays (black, blue, green and red balls) for each MgCl2 concentration. While the ternary complex with 0.5 or 1.0 mM MgCl2 did not cleave the DNA, the complex with 2.0 and 5.0 mM MgCl2 cleaved 39 ± 3% and 51  6% (mean ± SEM, n = 4) of the DNA, respectively. C, D FRET efficiency histograms of fluctuating (C) and static (D) S355C-S867C molecules. The panels from top to bottom show data in the presence of 0.5, 1 and 5 mM MgCl2. All of the assays were performed in the presence of 0.5 mM EDTA. The low, middle and high FRET efficiencies corresponding the I, D* and D positions of the HNH domain are indicated by black, green and blue arrowheads, respectively (C). The DNA cleavage activity (A, B) correlated well with the appearance of the high FRET efficiency peak (C), providing evidence that Cas9 molecules with the HNH domain in the D position are in the cleavage competent state.

**Figure EV5.**
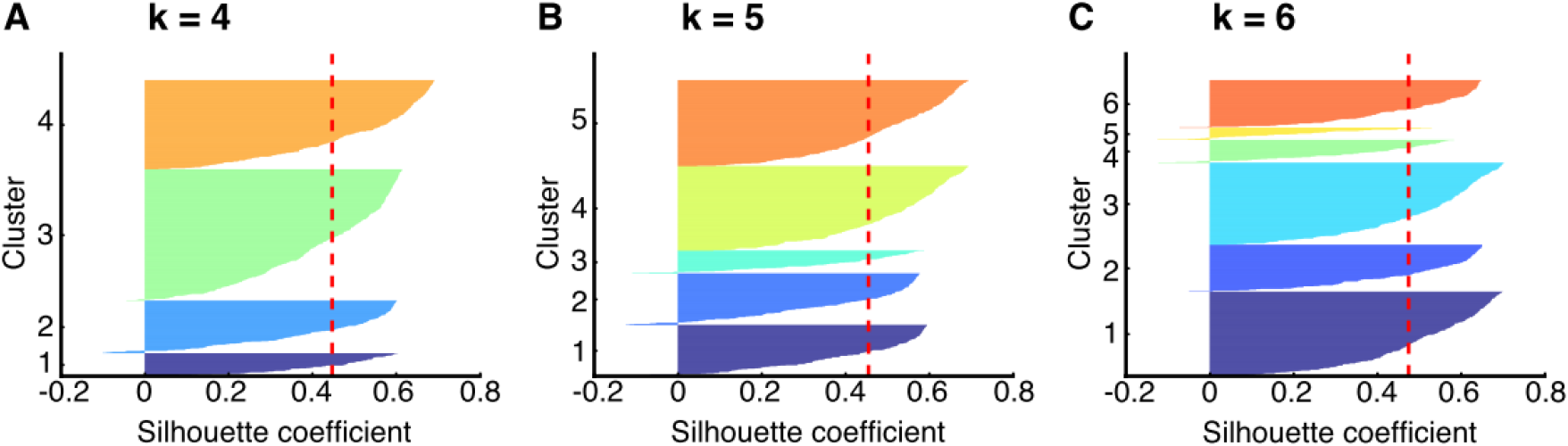
Silhouette analysis on k-means clustering of the FRET efficiency shift to determine the number of clusters. The Silhouette coefficients, which were calculated using the machine learning Python Package Scikit learn, were plotted for each cluster in the cases of k=4, 5 and 6, respectively (A-C). The vertical red dashed lines indicate the mean value of the Silhouette coefficients. In the cases of k=4 (A) and 5 (B), all clusters showed higher Silhouette coefficients than the mean values. This was not true for k=6, meaning that k=5 is the most probable number of clusters for the transition density plot shown in Fig 5B.

**Table EV1.** List of parameters for smFRET measurements.

|  |  | The distance of the dyes estimated from the crystal structure | The ratio of molecules showing FRET / total Cy3- and Cy5- doubly labeled molecules (%) | The number of analyzed fluorescent molecules (Figure 2 and 3D) | The number of traces showing fluctuations (Figure 4 and EV3) |
| --- | --- | --- | --- | --- | --- |
| **D435C-E945C (REC lobe-RuvC domain)** | No Nucleotide | 21 Å | 91 ± 2 | 121 | 14 |
|  | sgRNA | 78 Å | 90 ± 2 | 92 | 13 |
|  | sgRNA/DNA | 77 Å | 91 ± 2 | 98 | 26 |
| **S355C-S867C (REC lobe-HNH domain)** | No Nucleotide | 79 Å | 68 ± 2 | 191 | 79 |
|  | sgRNA | 81 Å | 79 ± 3 | 128 | 25 |
|  | sgRNA/DNA | 61 Å | 92 ± 2 | 140 | 42 |
| **S867C-N1054C (HNH domain-RuvC domain)** | No Nucleotide | 6 Å | 95 ± 2 | 103 | 4 |
|  | sgRNA | 7 Å | 96 ± 2 | 84 | 10 |
|  | sgRNA/DNA | 34 Å | 89 ± 2 | 107 | 6 |

## Appendix

**Appendix Figure S1.**
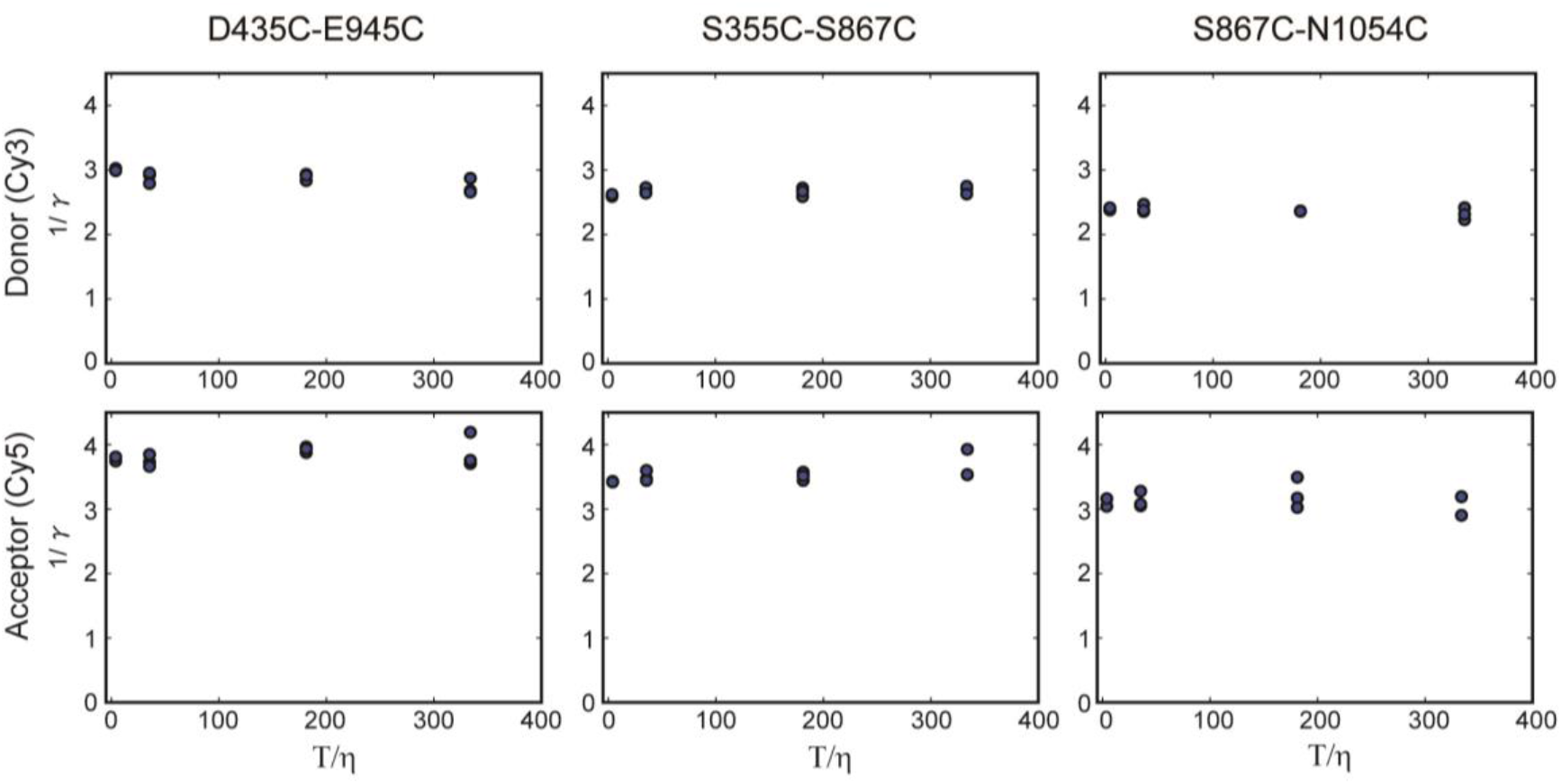
Perrin plots to calculate fluorescent anisotropy. The plots of the inverse of fluorescence anisotropies (*γ*) of Cy3 and Cy5 on the Cas9 constructs against *T*/*η*. Here, the absolute temperature *T* = 298 K, and the viscosities of the sample *η* were 0.89, 1.64, 8.39 and 75.89, corresponding to 0, 0.001, 0.01 and 0.1% methyl cellulose solutions, respectively. The plots are summaries of three individual experiments for each condition. The y-intercepts were calculated by extrapolating the plots to a linear function, yielding the estimated anisotropy values. The values of Cy3 anisotropy were 0.34 ± 0.006 in D435C-E945C, 0.38 ± 0.004 in S355C-S867C and 0.41 ± 0.004 in S867C-N1054C (mean ± SEM, n = 3). For Cy5, *γ* = 0.27 ± 0.005 in D435C-E945C, 0.29 ± 0.006 in S355C-S867C and 0.32 ± 0.009 in S867C-N1054C (mean ± SEM, n = 3). In the case of low anisotropy, the orientation factor *κ*^2^ is close to the dynamic isotropic limit of *κ*^2^ = 2/3. Otherwise, *κ*^2^ is widely distributed in the range of 0 ≤*κ*^2^≤4. Thus, the high anisotropies of Cy3 and Cy5 obtained here, which are close to the theoretical maximum value of 0.4, obscured the value of *κ*^2^, so that we were unable to estimate accurate distances between the two fluorochromes on the Cas9 molecules from the FRET efficiency.

**Appendix Figure S2.**
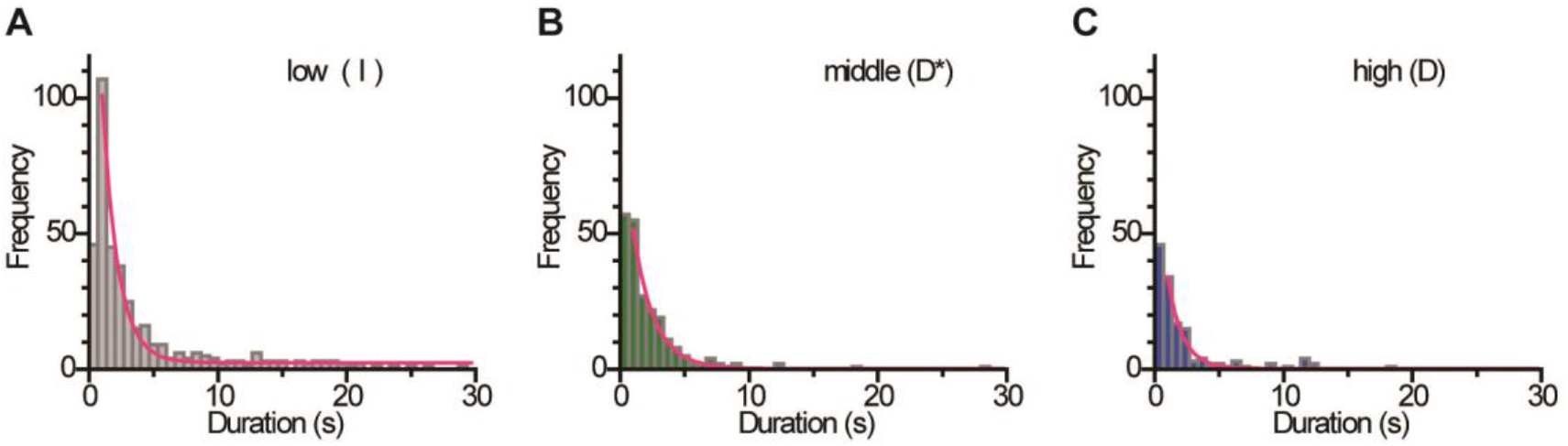
Dwell time histograms of the HNH domain in the three positions during flexible movements. A-C Dwell time distributions of the HNH domain in the I (A), D* (B) and D (C) positions in the fluctuating S355C-S867C molecules. The assays were performed in the presence of Mg^2+^, sgRNA and target DNA. By fitting the distributions to a single exponential decay function (red curves), the mean dwell times were determined as 1.22 ± 0.07 s for the I position (A: n = 399), 1.61 ± 0.08 s for the D* position (B: n = 219) and 1.14 ± 0.08 s for the D position (C: n = 124). Data: mean ± SEM.

**Appendix Figure S3.**
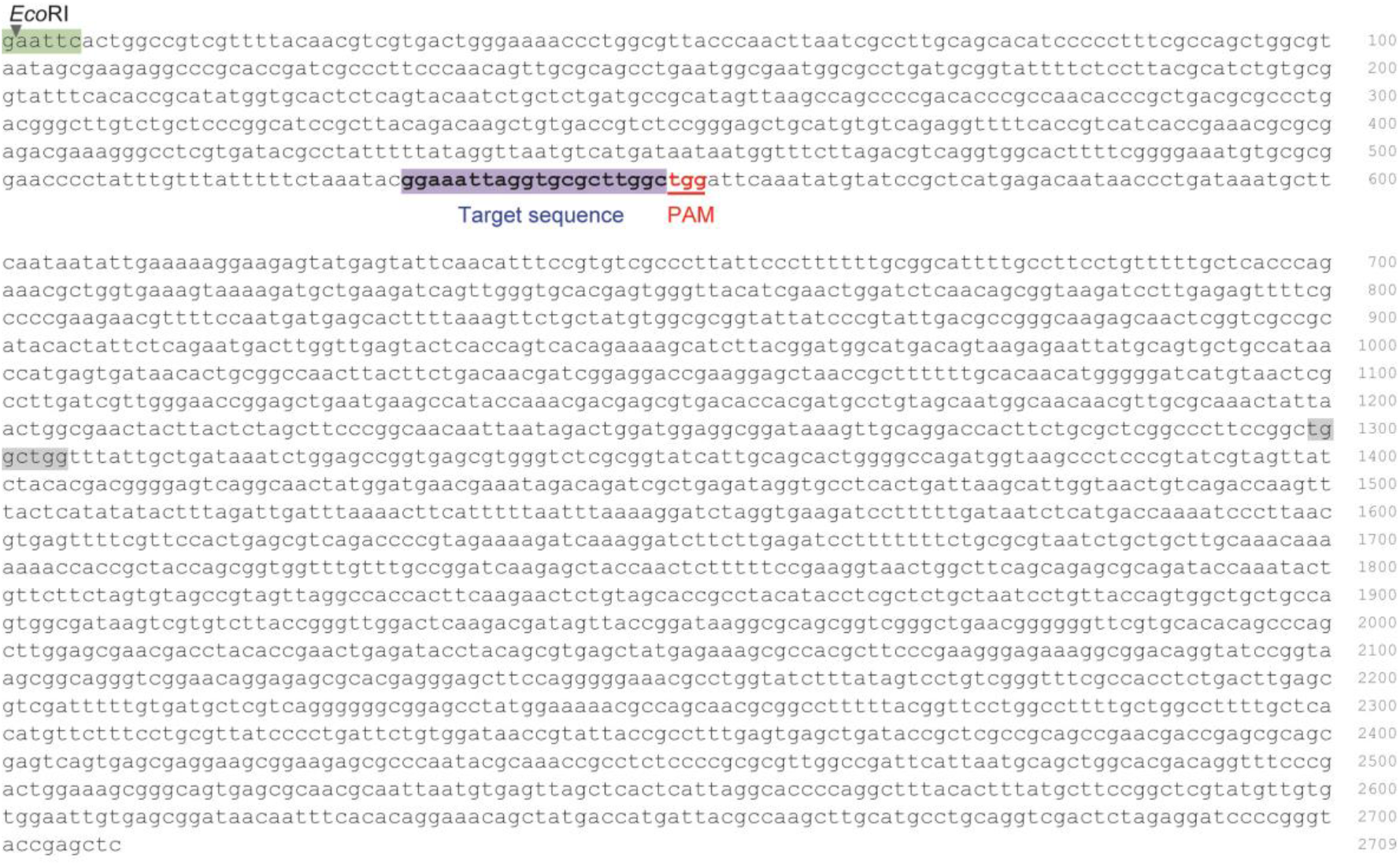
DNA sequence used in this study. The pUC119 plasmid containing the 20-nt target sequence (blue) and the NGG PAM (red) was linearized by *Eco*RI (green) and used as the target DNA. The longest off-target matching sequence to the sgRNA was 4-nt with a PAM sequence (grey highlight). Since the Cas9 binding to such a short matching sequence is highly unstable (Singh *et al*., 2016), we conclude that almost all of the Cas9 in the sgRNA/DNA-Cas9 ternary complex observed here was bound to the target sequence in the DNA.

